# Voxelated Bioprinting of Mechanically Robust Multiscale Porous Scaffolds for Pancreatic Islets

**DOI:** 10.1101/2021.06.29.449587

**Authors:** Jinchang Zhu, Yi He, Linlin Kong, Zhijian He, Kaylen Y. Kang, Shannon P. Grady, Leander Q. Nguyen, Dong Chen, Yong Wang, Jose Oberholzer, Li-Heng Cai

**Author notes:** Correspondence: L.-H. Cai, **Corresponding author contact:** Dr. Li-Heng Cai 228 Wilsdorf Hall, University of Virginia, 395 McCormick Road, Charlottesville, VA 22904. These authors contribute equally (Jinchang Zhu, Yi He).

## Abstract

Three-dimensional (3D) bioprinting additively assembles bio-inks to manufacture tissue-mimicking biological constructs, but with the typical building blocks limited to one-dimensional filaments. Here, we develop a technique for the digital assembly of spherical particles (DASP), which are effectively zero-dimensional voxels – the basic unit of 3D structures. We show that DASP enables on-demand generation, deposition, and assembly of viscoelastic bio-ink droplets. We establish a phase diagram that outlines the viscoelasticity of bio-inks required for printing spherical particles of good fidelity. Moreover, we develop a strategy for engineering bio-inks with independently controllable viscoelasticity and mesh size. Using DASP, we create mechanically robust, multiscale porous scaffolds composed of interconnected yet distinguishable hydrogel particles. Finally, we demonstrate the application of the scaffolds in encapsulating human pancreatic islets for responsive insulin release. Together with the knowledge of bio-ink design, DASP might be used to engineer highly heterogeneous yet tightly organized tissue constructs for therapeutic applications.

**Table of Contents:** A three-dimensional bioprinting technology is developed to enable on-demand generation, deposition, and assembly of viscoelastic bio-ink droplets in a biofriendly environment. A strategy is developed to independently control elasticity, viscosity, and mesh size of bio-inks. The technique allows for easy manufacturing mechanically robust multiscale porous scaffolds, which can be used to encapsulate human pancreatic islets for sustained responsive insulin release.

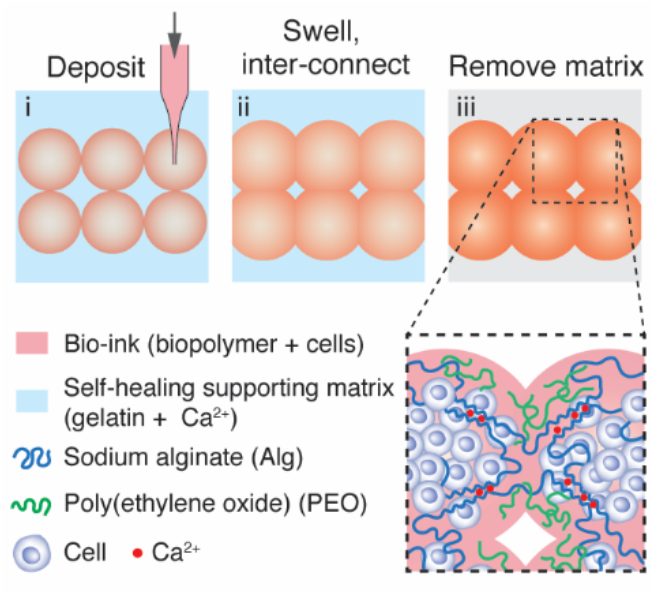

## Introduction

Three-dimensional (3D) bioprinting additively assembles bio-inks to manufacture tissue-mimicking biological constructs with broad applications in modeling tissue development and disease, drug screening, tissue engineering, and regenerative medicine ^[1,2]^. Unlike typical inks such as synthetic plastics and elastomers ^[3,4]^, bio-inks contain not only biopolymers but also living cells, which are sensitive to harsh processing conditions often involved in 3D printing; examples include UV light in stereolithography ^[5]^ and high mechanical shear in inkjet-based printing ^[6]^. These limitations can be circumvented by extrusion bioprinting, which exploits either pneumatic or mechanical pressure to physically extrude highly viscoelastic bio-inks without chemical interventions ^[7]^. Such mild processing conditions have negligible impact on the viability of cells even at a very high density comparable to that of organs ^[8]^. As a result, extrusion bioprinting becomes the most common technique for biofabrication ^[9,10]^.

The capability of extrusion bioprinting can be dramatically augmented by writing in a sacrificial supporting matrix ^[11–14]^, which is a yield-stress fluid ^[15]^ that reversibly transitions from solid-like to liquid-like at critical stress. During the process of printing, the supporting matrix entraps, holds, and mechanically supports the otherwise easily collapsed printed components; after printing, the sacrificial matrix can be removed to leave behind the printed 3D structures. This so-called embedded 3D bioprinting enables the fabrication of complex geometries and architectures mimicking that of major organs such as heart ^[16]^. Moreover, recently this technique has been extended to dispense liquid droplets for particle synthesis ^[17]^ and to fuse spheroids for high-cell density heterogeneous tissue models ^[18]^. Nevertheless, nearly all existing extrusion bioprinting platforms rely on layer-by-layer assembly of one-dimensional (1D) bio-ink filaments. This inevitably produces bulky, dense constructs with porosity determined by the mesh size of the bio-ink hydrogel. Unfortunately, the accessible range of the hydrogel mesh size is very limited ^[19]^, because the biopolymer must be above a certain concentration to achieve the viscosity required for printing and the stiffness required for maintaining the integrity of the printed structures ^[20–22]^. Introducing vascular networks ^[23–26]^ improves nutrient transport and waste exchange, which requires continuous external perfusion, however. This may limit the application of 3D printed biological constructs *in vivo*, where external perfusion is not available, and the transport of biomolecules must rely on diffusion. Moreover, the vascular networks are delicate and can be easily damaged by mechanical manipulation during transplantation, a process often involved in clinical applications. Consequently, it is highly needed a technique for manufacturing mechanically robust, multiscale porous scaffolds efficient for diffusion-based transport.

Here, we develop a 3D bioprinting technique that allows for the digital assembly of spherical particles (DASP), as schematically illustrated in **Figure 1** and with the hardware listed in Supporting Information (**SI Materials and Methods, Figure S1**). Unlike existing droplet-based 3D printing techniques that can disperse low viscosity fluids only, DASP enables on-demand generation, deposition, and assembly of viscoelastic bio-ink droplets in a biofriendly environment. This is accomplished through the development of *hybrid* bio-inks, which allows for independent control over elasticity, viscosity, and mesh size, three of the most important yet intrinsically coupled physical properties for biomaterials. The *hybrid* bio-ink is composed of two kinds of polymers interpenetrating with each other; one is an inert, long polymer that can form entanglements, whereas the other is a reactive polymer that can be crosslinked to form a network. The crosslinked polymer determines the network mesh size, whereas the entanglements determine the viscoelasticity of bio-inks. Using DASP, we create scaffolds composed of interconnected yet distinguishable hydrogel particles. We show that these scaffolds are not only mechanically robust but also of multiscale porosity: sub-millimeter pores determined by the interstitial space among particles, and sub-micrometer pores determined by the mesh size of the hydrogel network. Finally, we demonstrate the application of the scaffolds in encapsulating human pancreatic islets, and that the multiscale porous nature enables sustained, responsive insulin release in response to glucose stimulation. Considering that spherical particles are effectively zero-dimensional (0D) voxels – the basic building blocks of 3D structures, DASP might be used to engineer highly heterogeneous yet organized 3D tissue constructs for fundamental biomedical research and therapeutic applications.

**Figure 1.**
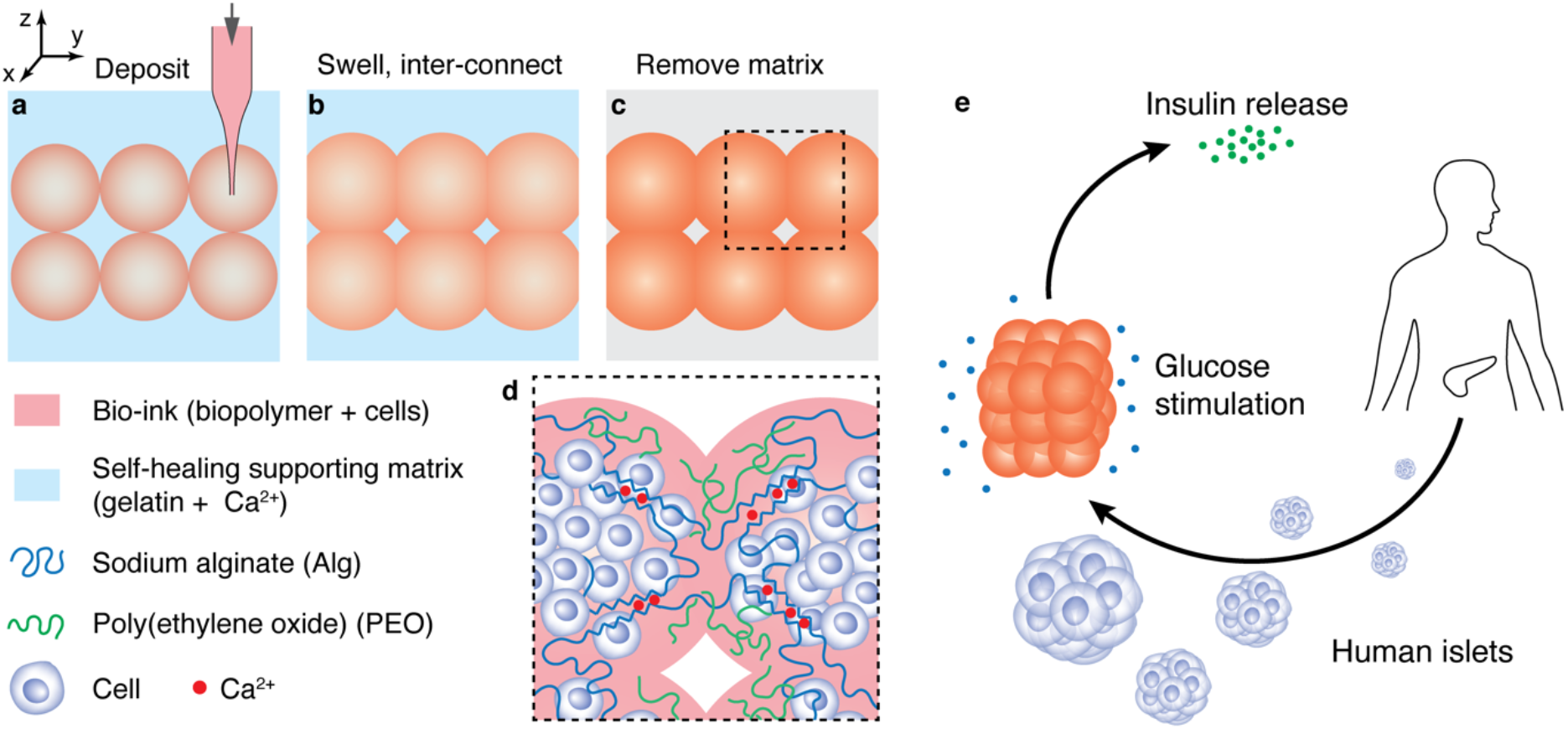
Concept of digital assembly of spherical particles (DASP). a) Bio-ink particles are deposited at precisely controlled locations and with a prescribed volume in a sacrificial supporting matrix. The bio-ink is a mixture of cells and a solution consisting of two polymers, alginate and poly(ethylene oxide) (PEO). The supporting matrix is a slurry of densely packed gelatin microparticles and contains calcium ions. b) A bio-ink particle swells to partially coalesce with its neighbors. In the meantime, calcium ions in the supporting matrix diffuse into the bio-ink particle to partially crosslink the alginate. c) Following complete crosslinking of the alginate network and subsequent removal of the sacrificial supporting matrix, the printed particles form a 3D lattice of interconnected yet distinguishable hydrogel particles. d) The resulted hydrogel scaffold is of multiscale porosity: sub-millimeter pores determined by the interstitial space among particles, and sub-micrometer pores determined by the mesh size of the hydrogel network. e) The workflow of using DASP to encapsulate human islets for responsive insulin release.

## Results and Discussion

### Design and development of the supporting matrix

A suitable supporting matrix for DASP must possess five features. First, the matrix must be aqueous and cytocompatible. Second, the matrix must allow the printing nozzle to move through without mechanically perturbing the printed structures (**Figure 1a**). Third, once a particle is deposited and the printing nozzle leaves its original position, the matrix must be able to self-heal to recover its original mechanical properties within a short time to localize the particle ^[14]^. Fourth, the supporting matrix must provide an environment in which individual particles can swell and to partially coalesce with each other, the mechanism by which particles interconnect with each other to form an integrated 3D structure (**Figure 1b**). Finally, the matrix must be easily removed after printing (**Figure 1c**).

We extend a recently developed concept that relies on shear-induced unjamming of microparticles ^[12,13]^ to engineer such a supporting matrix. The matrix is a slurry of fragmented hydrogel microparticles made of gelatin, a biocompatible material widely used in tissue engineering. Therefore, by design the matrix is cytocompatible and easily removable. By exploring different processing conditions (**SI Materials and Methods**), we identify a gelatin matrix that is solid-like with a shear storage modulus, *G*′ ≈ 130 *Pa*, significantly larger than the loss modulus, *G″* ≈ 7 *Pa*, corresponding to a small loss factor tan *δ ≡ G*^*″*^/*G*^′^ ≈ 0.05 (circles in **Figure 2a**). Moreover, the matrix yields above a critical shear stress of 10 Pa, near which *G*′ decreases dramatically to ∼1 Pa and becomes much lower than *G″*. Simultaneously, the shear strain exhibits a sudden increase from 10% to 1000%, suggesting that the matrix flows (the squares in **Figure 2a**). These results demonstrate that the gelatin matrix is a yield-stress fluid, which transitions from solid-like to liquid-like above the critical shear stress. This feature allows the printing nozzle to move freely through the supporting matrix.

**Figure 2.**
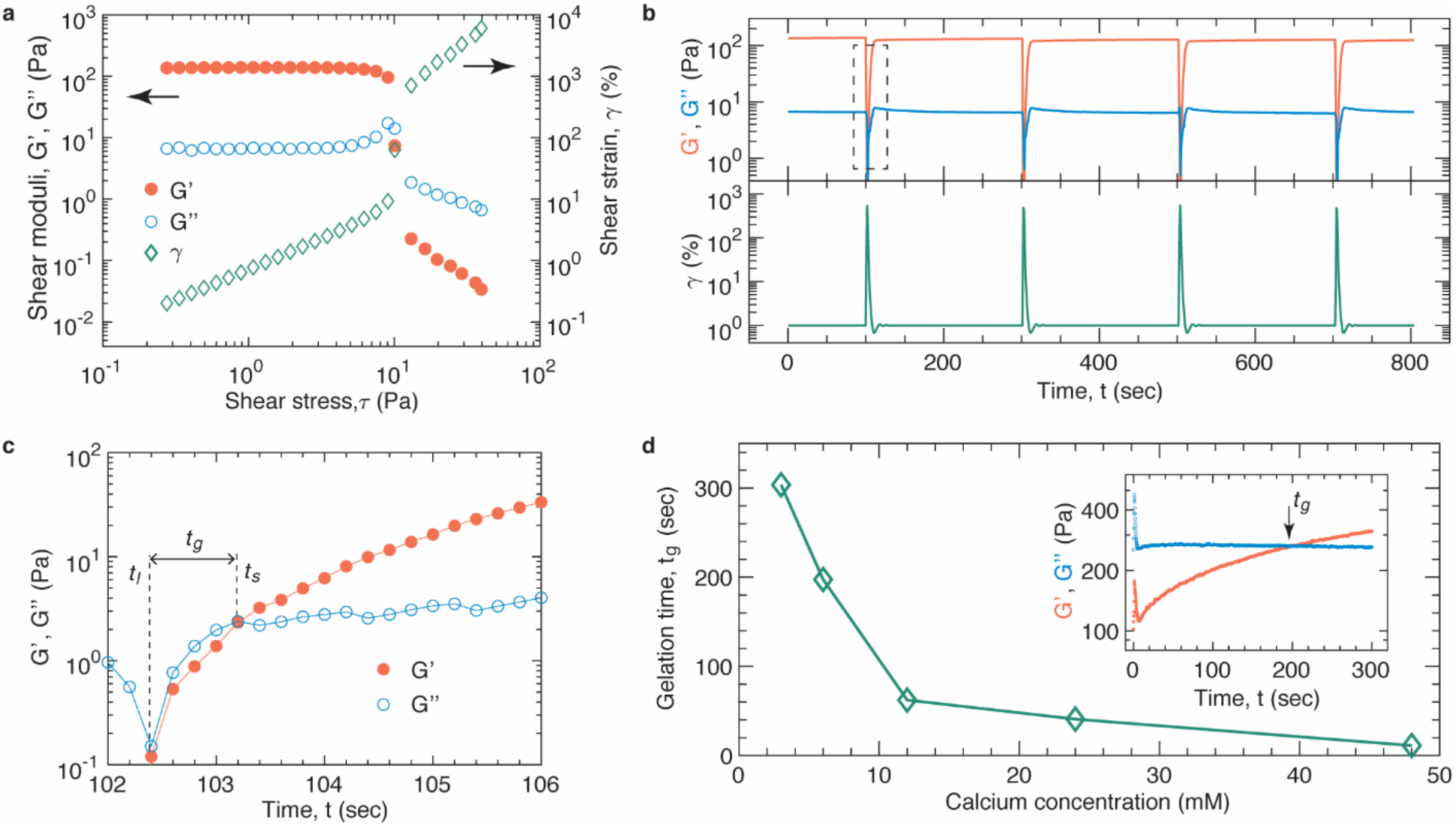
Physical and chemical properties of the gelatin supporting matrix and the alginate hydrogel. a) The supporting gelatin matrix is a yield-stress fluid that becomes fluid-like at shear stresses above 10 Pa. Left *y*-axis: the dependence of storage modulus *G*′ (filled circles), and loss modulus *G″* (empty circles), on shear stress *τ*. Right y-axis: the dependence of shear strain (empty squares), *γ*, on shear stress. The measurements are performed at oscillatory frequency of 1 Hz at 20 *°*C. b) The supporting matrix self-heals to recover its original properties for unlimited times. Upper: *Real-time* behavior of *G*′ (red symbols) and *G″* (blue symbols) under a periodic destructive high shear strain of 1000%. The high shear strain is applied for 1 sec, and the waiting time between two neighboring destructive shear strains is 200 seconds. c) After the shear induced fluidization at *t*_*l*_, the supporting matrix becomes solid like at time scales *t > t*_*s*_, at which *G*′ is larger than *G″*. The gelation time, *t*_*g*_ = *t*_*s*_ *– t*_*l*_, is less than 1 sec. d) Crosslinking kinetics of the alginate hydrogel: Dependence of the gelation time on the concentration of calcium ions. Insert: An example *real-time* characterization of *G*′ (red symbols) and *G″* (blue symbols) for the alginate solution following its contact with the gelatin supporting matrix that contains 6 mM Ca^2+^. The peaks for *G*′ and *G″* are associated with the instantaneous increase in friction between the oscillatory geometry and the gelatin supporting matrix.

To test whether the fluidized gelatin matrix can self-heal, we apply a large, instant shear of 1000% strain to break the matrix, and simultaneously monitor in *real-time* the recovery of mechanical properties. The gelatin matrix completely reverts to its original mechanical properties, and this damage-recovery process is repeatable, as shown in **Figure 2b**. Moreover, after a destructive shear, the fluidized gelatin matrix becomes solid-like again within less than 1 second, as shown by gelation kinetics in **Figure 2c**. These results indicate that the gelatin matrix can self-heal in a short time. Moreover, the matrix is semi-transparent to permit direct visualization of printed structures. These features make the material an ideal supporting matrix for embedded 3D bioprinting.

To make the gelatin matrix suitable for DASP, we introduce Ca^2+^ ions into the matrix for crosslinking alginate-based particles. Yet, the time for a particle to solidify must be long enough for printing multiple particles and for individual hydrogel particles to swell to partially coalesce with neighbors. Therefore, the crosslinking kinetics for individual particles must be carefully controlled to ensure a balance between the particle solidification and the printing speed. To this end, we quantify the crosslinking kinetics of alginate at various concentrations of calcium ions. We monitor in *real-time* the viscoelasticity of the alginate solution exposed to calcium contained gelatin matrix using a stress-controlled rheometer (**SI Materials and Methods**). This allows us to determine the gelation time of alginate, *t*_*g*_, above which *G*′ becomes larger than *G″*, as illustrated by the insert in **Figure 2d**. The gelation time decreases by nearly 30 times from 300 to 10 seconds as the Ca^2+^ concentration increases from 3 to 48 mM (**Figure 2d** and **Figure S2a-e**). Comparing the gelation time to the swelling kinetics of an alginate particle, which takes about 200 seconds to reach 80% of equilibrium swelling (**Figure S3**), we determine the optimum Ca^2+^ concentration 6 mM associated with a gelation time of 200 seconds. This time is long enough for two neighboring particles to partially coalesce before solidified. Moreover, comparing to the printing speed, which takes about 3 seconds to print a particle (**Movie S1**), this time allows for printing tens of hydrogel particles before they are partially solidified. Taken together, these studies establish the design parameters and provide a formulation for the sacrificial supporting matrix required for DASP.

### Design strategy of bio-inks for DASP

We design bio-inks based on two criteria: (1) 3D printability and (2) porosity of the crosslinked hydrogel particles. To identify the physical parameters that determine the 3D printability of bio-inks, we consider the whole process for printing a hydrogel particle. The process involves three steps: (i) deposit an alginate-based particle with a prescribed volume in the supporting matrix; (ii) detach the printing nozzle, or the glass capillary tip, from the particle; (iii) wait for the particle to be partially crosslinked. Unlike depositing an aqueous droplet in an oil-based yield-stress fluid ^[17]^, where the droplet spontaneously adopts a spherical shape to minimize the interfacial free energy between the two immiscible fluids, in our case, both the bio-ink particle and the supporting matrix are aqueous and have negligible interfacial tension. Moreover, although being optically semi-transparent, the supporting matrix is inhomogeneous at microscopic scale, because it consists of fragmented gelatin microparticles with a wide distribution in size and shape, as evidenced by previous studies ^[12]^ and illustrated in **Figure 3a**. To maintain good fidelity, therefore, the bio-ink must have a relatively high viscosity such that a printed particle does not flow through the porous supporting matrix in a short time. In addition, the alginate-based bio-ink is not a simple Newtonian but a complex viscoelastic fluid. This poses a challenge for particle printing, during which the dragging force associated with detaching the glass capillary from the fluid may mechanically deform the particle. And such a dragging effect is expected to be more pronounced for a more elastic particle. As a result, to be suitable for particle printing, we postulate that the bio-ink must have relatively a high viscosity yet a low elasticity, or a high loss factor, as illustrated in **Figure. 3b**.

**Figure 3.**
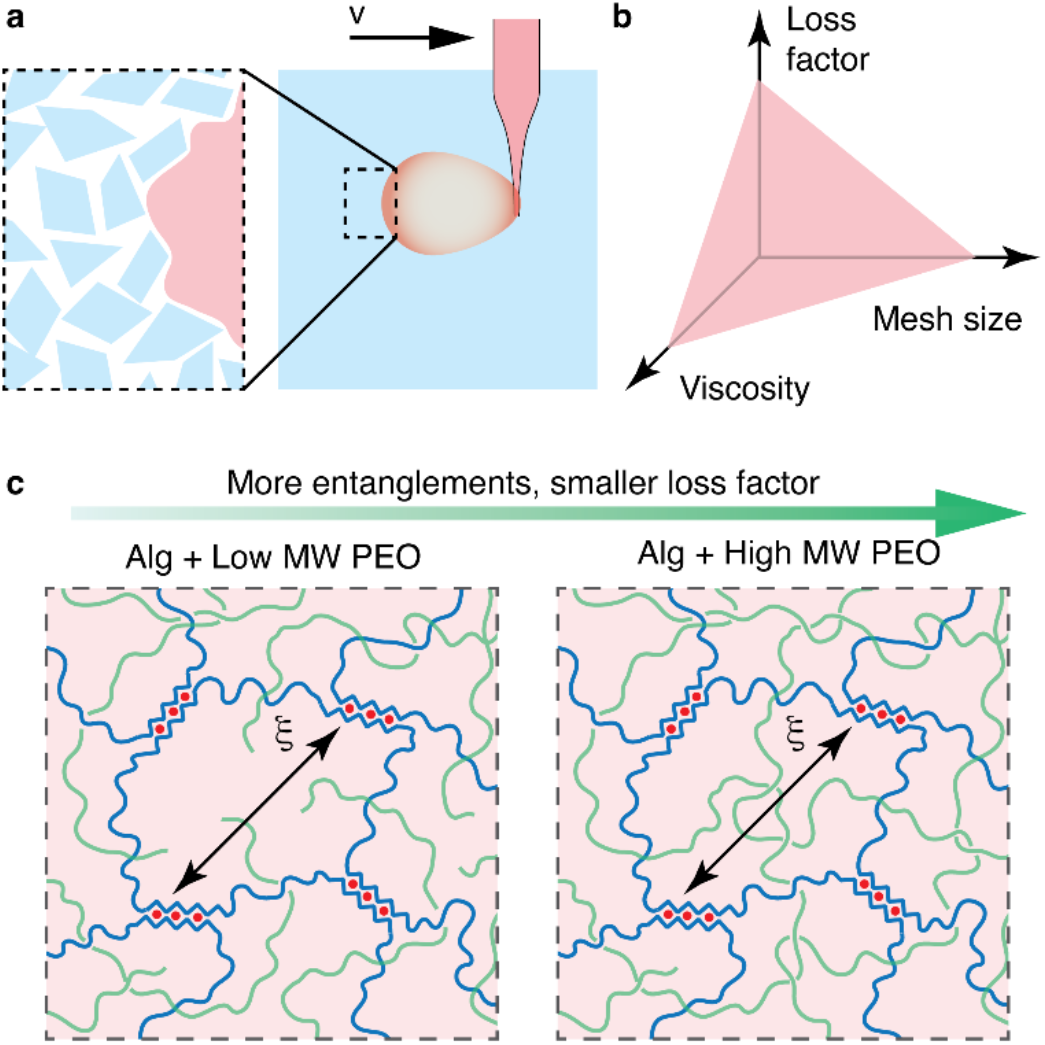
Bio-ink design. a) Illustration of the process of printing a bio-ink particle: (i) deposit an alginate-based particle with a prescribed volume in the supporting matrix; (ii) detach the glass capillary tip from the particle; (iii) wait for the particle to be partially crosslinked. The supporting matrix is a heterogeneous, porous environment filled with gelatin microparticles of irregular shape. b) To be suitable for DASP, a bio-ink must have optimized viscosity, loss factor, and mesh size. c) A *hybrid* bio-ink exploits crosslinked alginate to control mesh size and entangled PEO polymers to control viscosity and loss factor.

Besides viscoelasticity, the porosity of the bio-inks must be tunable to meet the requirement of various applications ^[17]^. For example, efficient nutrient transport requires a relatively large hydrogel mesh size. By contrast, in some applications such as encapsulating islets to treat type 1 diabetes, hydrogels with a relatively tight mesh are needed to protect encapsulated cells from immune attack ^[27]^. Unfortunately, the mesh size and the viscosity of a hydrogel are intrinsically coupled as they both increase with the polymer concentration. To solve this challenge, we create an *hybrid* bio-ink consisting of alginate and poly(ethylene oxide) (PEO), a polymer widely used for biomedical applications ^[28]^. In the *hybrid* bio-ink, the alginate polymer can form a crosslinked network that determines the hydrogel mesh size. By contrast, the long linear PEO polymers do not crosslink but form physical entanglements, which are known to be the dominating factor responsible for the viscoelasticity of polymer solutions ^[29]^. As a result, the concept of *hybrid* bio-inks is expected to enable independent control over the mesh size and the viscoelasticity, as illustrated in **Figure 3c**.

### Development of bio-inks for DASP

Guided by above design strategy, we first determine the bio-ink viscoelasticity required for DASP. To do so, we print pure alginate particles of various concentrations and examine printing fidelity. We start with alginate concentration of 1.5% (w/v) (Alg^1.5^), a formulation widely used for cell encapsulation and biomaterial design because of balanced mechanical properties and mesh size ^[30]^. Moreover, a fluorescently labeled alginate is used to allow direct visualization of the particles by optical microscopy (**SI Materials and Methods**). The printed particle of Alg^1.5^ exhibits an irregular shape, as shown by the optical image on the top left in **Figure 4**a. Increasing alginate concentration improves the printing fidelity, but only at concentrations no lower than 4.0% (w/v) the particles become spherical (top panel, **Figure 4a**). These results indicate that the minimum concentration required for printing an alginate particle with good fidelity is 4.0% (w/v) (Alg^4.0^). The viscosity of Alg^4.0^ is about 130 Pa·s, which is about 30 times of the viscosity of Alg^1.5^, as shown respectively by the empty and filled red symbols in **Figure 4b**. Moreover, Alg^4.0^ is of a relatively low elasticity, as reflected by a high loss factor, tan δ *≡ G*^*″*^/*G*^′^ = 1.56 (**Figure 4c**,**d**). These studies identify the viscosity and the loss factor required for printing an alginate particle of good fidelity.

**Figure 4.**
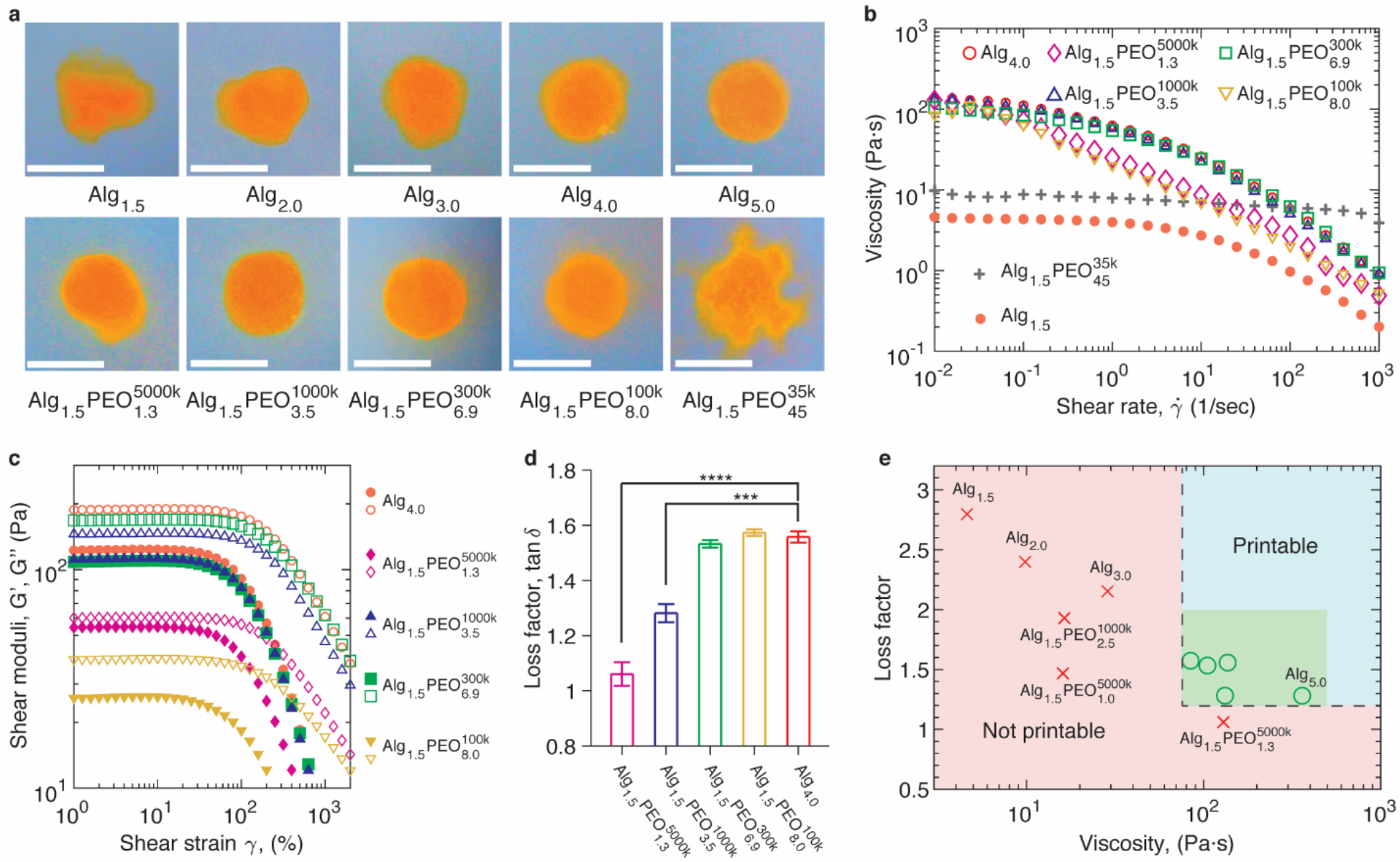
Rheological properties of bio-inks required for printing hydrogel particles with a good fidelity. ) Optical images of fluorescently labeled hydrogel particles printed at the same conditions but using bio-inks of different compositions. A bio-ink is denoted as Alg_*x*_PEO_*y*_^*w*^, in which *x* and *y* are respectively the concentration (w/v) of alginate (Alg) and poly(ethylene oxide) (PEO), and *w* is the molecular weight (MW) of PEO in g/mol. Scale bars, 1 mm. b) By adjusting the MW and the concentration of PEO, the viscosities of Alg/PEO hybrid bio-inks can be tuned to be close to that of pure alginate Alg_4.0_, which has the minimum concentration required for printing spherical hydrogel particles with good fidelity. c) Large amplitude oscillatory shear measurements for composites bio-inks with viscosities matching that of Alg_4.0_. Solid symbols, storage modulus *G*′; empty symbols, loss modulus *G″*. Measurements are performed at 20 °C and 1 Hz. d) The loss factor (tanδ ≡ *G″*/*G*′) bio-inks in (c). Error bar, standard deviation for *n=3*. \*\*\**p*<0.001, \*\*\*\**p*<0.0001 e) A two-parameter (viscosity and loss factor) phase diagram that outlines the rheological properties required for printing hydrogel particles with good fidelity. Green area denotes the viscoelasticity that can be easily achieved for bio-inks. Bio-inks in the light blue region should be printable but relatively challenging to design because of high viscosity and loss factor.

Next, we engineer alginate/PEO *hybrid* bio-inks with viscoelasticity matching that of Alg^4.0^. We fix the alginate concentration at 1.5% (w/v) while changing the concentration of PEO polymers. For the PEO of a high molecular weight (MW) 5000 kDa, a small addition at the concentration of 1.3% (w/v) results in a hybrid bio-ink, Alg^1.5^PEO_1.3_^5000k^, of a viscosity comparable to that of Alg_4.0_, as shown by the empty diamonds in **Figure 4b**. However, the loss factor of Alg_1.5_PEO_1.3_^5000k^ is 1.06±0.04, significantly smaller than the value 1.56±0.02 for Alg_4.0_ (*p*<0.0001) (diamonds in **Figure 4c** and pink bar in **Figure 4d**). This is likely because the PEO polymers are too long to relax at the probed time scale, such that the entanglements effectively act as crosslinks to result in a relatively elastic bio-ink. Compared to a viscous particle with a high loss factor, an elastic particle with a low loss factor is more likely to be deformed by the dragging force associated with detaching the printing nozzle from the particle. Consistent with this understanding, the printed particle Alg_1.5_PEO_1.3_^5000k^ exhibits a distorted morphology (lower left of **Figure 4a**).

To create bio-inks with high loss factors, we lower the MW but increase the concentration of PEO polymers. As the PEO MW decreases from 5000 kDa to 100 kDa, to match the viscosity of Alg^4.0^, the PEO concentration must increase by nearly 6 times from 1.3% (w/v) to 8.0% (w/v); concurrently, the loss factor increases from 1.06 to 1.57. Using this strategy, we create three bio-inks, Alg^1.5^PEO_3.5_^1000k^, Alg_1.5_PEO_6.9_^300k^, and Alg_1.5_PEO_8.0_^100k^, with both the viscosity and the loss factor matching those of Alg_4.0_ (**Figure 4b-d**). All these three formulations can be used to print particles of good fidelity (lower panel, **Figure 4a**). By contrast, further decreasing the PEO MW to 35 kDa does not permit a bio-ink with viscosity comparable to Alg_4.0_ even at a very high PEO concentration 45% (w/v) (Alg_1.5_PEO_45_^35k^, down-pointing triangles, **Figure 4b**). This is likely because the MW is too low to form highly entangled polymers, which is required to achieve a high viscosity. Because of the low viscosity, the bio-ink flows through the porous supporting matrix, as reflected by the multiple protrusions of a particle as shown by the lower right image in **Figure 4a**. Taken together, these studies validate our hypothesis that viscosity and loss factor are the two major parameters determining the printability of bio-inks. Based on these results, we construct a two-parameter phase diagram that delineates bio-ink formulations suitable for DASP, as outlined by the dashed lines in **Figure 4e**.

### Porosity of bio-ink hydrogel particles

The porosity of hydrogel particles determines their ability to exchange molecules between interior and external environments, and it is a critical parameter for biomaterial design ^[19]^. To characterize the porosity of a bio-ink hydrogel particle, we quantify the release profile of encapsulated molecules using time-lapsed fluorescence confocal microscopy (**SI Materials and Methods**). We use fluorescently labeled dextran with a MW of 70 kDa as the probe molecule. The dextran is an inert, randomly branched molecule with a weight average hydrodynamic radius of ∼7 nm, larger than the size of metabolically important biomolecules ^[31]^. For pure alginate hydrogel particles, as the alginate concentration increases from 1.5% (w/v) to 4.0% (w/v), the release of dextran molecules becomes noticeably prolonged, as shown by the time-lapse fluorescence images in **Figure 5a**. Quantitatively, the half-decay time of the fluorescence increases by nearly twice from 3.5 to 7 minutes (dash line in **Figure 5b**). By contrast, fixing the alginate concentration at 1.5% (w/v) while introducing PEO molecules results in a negligible increase of the release profile, as shown by the overlap between the profiles for Alg^1.5^ (circles) and Alg^1.5^PEO_8.0_^100k^ (triangles) in **Figure 5b**. These results provide direct evidence that the mesh size of a bio-ink hydrogel particle is determined by the concentration of the crosslinked alginate but not so much by the uncrosslinked PEO molecules.

**Figure 5.**
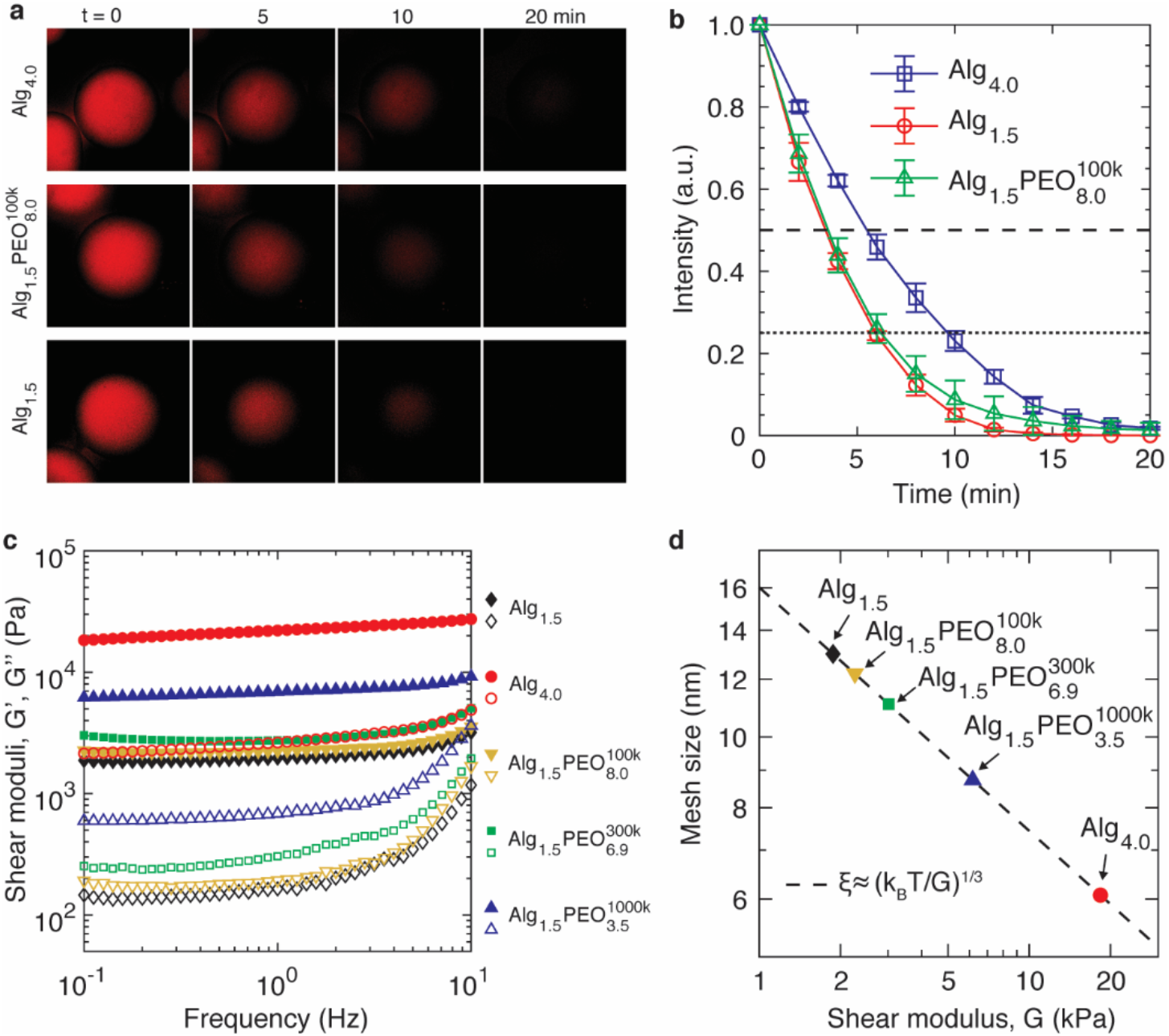
Porosity of crosslinked bio-ink particles. a) Time-lapse fluorescence confocal microscopy images showing the release of encapsulated fluorescently labeled dextran molecules (Texas Red, 70 kDa) from individual hydrogel particles formed by different bio-ink formulations. b) Dependence of the intensity on waiting time. The error bar is the standard deviation based on measurements from different batches of particles (*n=3*). c) Dynamic mechanical properties of crosslinked bio-ink hydrogels in the oscillatory shear frequency range of 0.1-10 Hz. Filled symbols are storage modulus, and empty ones are loss modulus. d) Estimated mesh size ξ based on the equilibrium shear storage modulus of bio-inks.

The difference in release profiles of bio-ink formulations is further confirmed by the variation in hydrogel stiffness. The equilibrium shear modulus of an unentangled polymer network is about *k*_*B*_*T* per volume of a network strand. Therefore, the network strand size, or mesh size, can be estimated based on the network shear modulus: *ξ* ≈ (*k*_*B*_*T*/*G*)^1/3^, where *k*_*B*_ is Boltzmann constant, and *T* is the absolute temperature ^[29]^. For the bio-ink formulations suitable for DASP, the shear storage moduli *G*^′^ are nearly independent of oscillatory shear frequency within the range of 0.1-10 Hz (filled symbols, **Figure 5c**); moreover, they are nearly 10 times larger than the loss moduli *G*^*″*^ (empty symbols, **Figure 5c**). These results suggest that all crosslinked hydrogels are elastic networks, such that we can take the values of *G*^′^ at the lowest frequency as the equilibrium shear moduli.

Interestingly, adding 100 kDa PEO polymers only increases the modulus slightly from 1.8 kPa to 2.2 kPa, which respectively correspond to bio-inks Alg_1.5_ and Alg_1.5_PEO_8.0_^100k^. However, using PEO molecules of higher MW, 1000 kDa, results in a larger increase in modulus to 6 kPa, as shown by the solid upward-triangles in **Figure 5c**. This is likely because the long PEO molecules are trapped in the alginate network and have not yet fully relaxed at the probing time scale 10 seconds, resulting in an effective increase in shear modulus. As a result, exploiting entanglements formed by long PEO molecules permits fine tuning the effective network mesh size in the range of 9 to 12 nm (**Figure 5d**). Yet, the mesh sizes of all crosslinked *hybrid* bio-ink hydrogels are much larger than 6 nm for pure alginate Alg_4.0_. Moreover, the trapped long PEO polymers are not bounded to the alginate network; instead, they can escape from the network through reptation, a mechanism for entangled polymers to diffuse at long time scales ^[29]^. Collectively, our results demonstrate that the concept of *hybrid* bio-inks enables independent control over the mesh size and the viscoelasticity.

### Implementation of DASP

Individual hydrogel particles are the building blocks for DASP. Therefore, it is critical to determine the relation between printing parameters and particle size. To this end, we print individual spherical particles at various combinations of injection speed and volume. We vary the injection speed by nearly 17 times from 40 to 680 nL/sec, and for each speed, we vary the injection volume by 27 times from 16 to 418 nL. This allows us to control the diameter of hydrogel particles from 300 μm to 1.2 mm, the size range suitable for encapsulating cells and pancreatic islets ^[32]^. Moreover, regardless of the injection speed, the particle diameter, *d*, scales with the injection volume *V* by *d*∼*V*^1/3^, as shown by the symbols in **Figure 6a**. This suggests that the particle size is determined by the injection volume but not so much by the injection speed.

**Figure 6.**
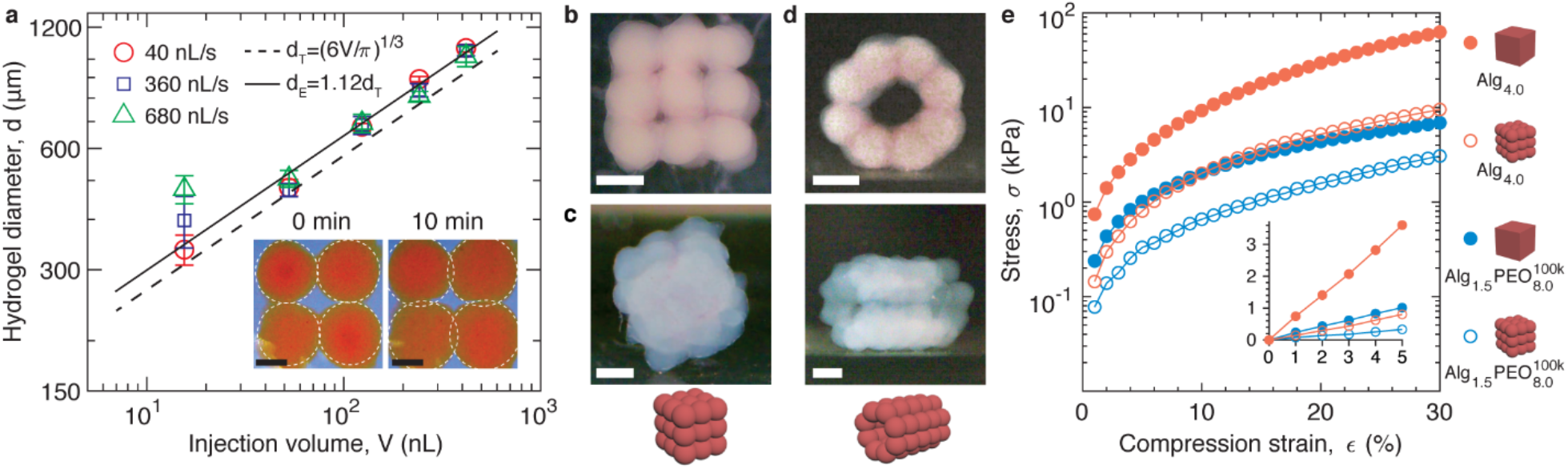
Implementation of DASP in printing mechanically robust, multiscale porous 3D structures. a) Dependence of hydrogel particle diameter on various combinations of injection volume and speed. Dashed line: predicted particle diameter *d*_*T*_ based on the injection volume *V*: *d*_*T*_ = (*6V*/*π*)^1/3^. Solid line is the best weighted fit to the experimental data: *d*_*E*_ = (1.12 ± 0.04)*d*_*T*_, suggesting the particles swell by 12%. Inset: Hydrogel particles swell by nearly 11% in diameter at 10 mins. A particle reaches 80% of equilibrium swelling at about 200 seconds (**Figure S3**), comparable to the gelation time of alginate (**Figure 2d**). Scale bars, 0.5 mm. Error bar, standard deviation for *n=3*. b-d) Printing a 3×3×3 lattice and a tube composed of 40 hydrogel particles made of bio-inks Alg_1.5_PEO_8.0_^100k^ (b) and Alg_4.0_ (c, d). In all structures, the particles are interconnected yet distinguishable. Scale bars, 1 mm. e) Mechanical behavior of printed 3D structures under compression.

Interestingly, the measured particle diameter is higher than predicted, and the difference between the two is higher for larger particles (dashed line in **Figure 6a**). Yet, the swelling ratio, defined as the experimentally measured particle diameter over the theoretical value, is nearly a constant of 1.12±0.04 for all particle sizes (solid line in **Figure 6a**). Consistent with this, direct visualization of fluorescently labeled alginate hydrogel particles reveals that a particle swells by nearly 11% in diameter (inset in **Figure 6a and Figure S3**).

We exploit the hydrogel swelling to partially coalesce two neighboring particles; this allows for creating 3D structures composed of interconnected yet distinguishable spherical hydrogel particles (**SI Materials and Methods**, and **Movie S1**). We start with printing a 3×3×3 lattice composed of hydrogel particles of 1.0 mm in diameter. The printed structure successfully recapitulates the design, in that the hydrogel particles are interconnected yet distinguishable, and this fidelity applies to both *hybrid* bio-inks (**Figure 6b**) and pure alginate (**Figure 6c**). Built on this success, we seek to print a 1D tube with the wall composed of a single layer of hydrogel particles. Contrasting to the flat surface in a 3D cuboid, the surface of the 1D tube is highly curved, which precludes the feasibility of the classical layer-by-layer assembly of bio-inks. Moreover, the tube wall has only one layer of particles, which can easily fall apart if a particle does not swell as expected or is off its prescribed position during the whole process of printing. Thus, printing such a 1D tube represents the ultimate test for the robustness and reliability of DASP. We successfully print a 1D tube with the wall composed of a single layer of 40 hydrogel particles. Remarkably, in such a tube, each particle is interconnected with each other yet distinguishable, precisely capturing the 3D design (**Figure 6d**). Collectively, these results demonstrate that DASP allows for facile, robust 3D digital assembly of highly viscoelastic hydrogel particles.

Unlike existing droplet assembly that relies on sophisticated surface chemistry such as lipid bilayers to bridge two neighboring particles ^[33,34]^, in DASP individual hydrogel particles are connected through crosslinked alginate polymer network. Therefore, although being porous because of the interstitial space among particles, the printed 3D structures are expected to be mechanically robust. To test this, we perform compression tests for a 3×3×3 lattice structure and compare its behavior to that of the bulk with the same dimension (**SI Materials and Methods**).

The compression modulus of the DASP printed porous lattice structure is about three times smaller than the bulk counterpart, and this difference applies to bio-inks of different stiffnesses, as shown by the red circles for Alg_1.5_PEO_8.0_^100k^ and by the blue circles for Alg_4.0_ in the inset of **Figure 6e**. Moreover, regardless of the bio-ink stiffness, the lattice structure can sustain a large compression strain of nearly 30% without fracture, as shown by open circles in **Figure 6e**. Consistent with the mechanical robustness, the lattice structure can be flipped, rotated, and translated without breaking apart (**Movie S2**). Taken together, our results demonstrate that DASP enables manufacturing free-standing, mechanically robust, multiscale-porous 3D structures.

### Cytocompatibility and Insulin Release

To demonstrate the potential of DASP in biomedical applications, we explore the DASP printed scaffolds in encapsulating pancreatic islets, which represents an emerging biophysical approach to treat type 1 diabetes by protecting islets from immune attack ^[32,35,36]^. We start with testing the cytocompatibility of bio-inks by using them to culture MIN6 cells, a pancreatic beta cell line that retains glucose-stimulated insulin secretion as isolated islets. Live/dead assay reveals that bio-inks made of Alg_4.0_ and Alg_1.5_PEO_8.0_^100k^ respectively exhibit a viability of 95.3±1.5% and 90.4±0.5% (**Figure S4**). Further, we test whether the mechanical shear involved in DASP impairs cell viability (**SI Materials and Methods**). We find that DASP encapsulated MIN6 cells maintains 95% viability, nearly the same as that of naked cells; moreover, such a remarkable viability remains for 3 days, a typical waiting time before transplantation in clinical settings (**Figure 7a, b**). Moreover, DASP encapsulated mice islets remain a very high viability, >80%, in 3 days, which has no significant difference compared to that of naked mice islets (**Figure 7c** and blue bars in **Figure 7e**). These results demonstrate that DASP is cytocompatible.

**Figure 7.**
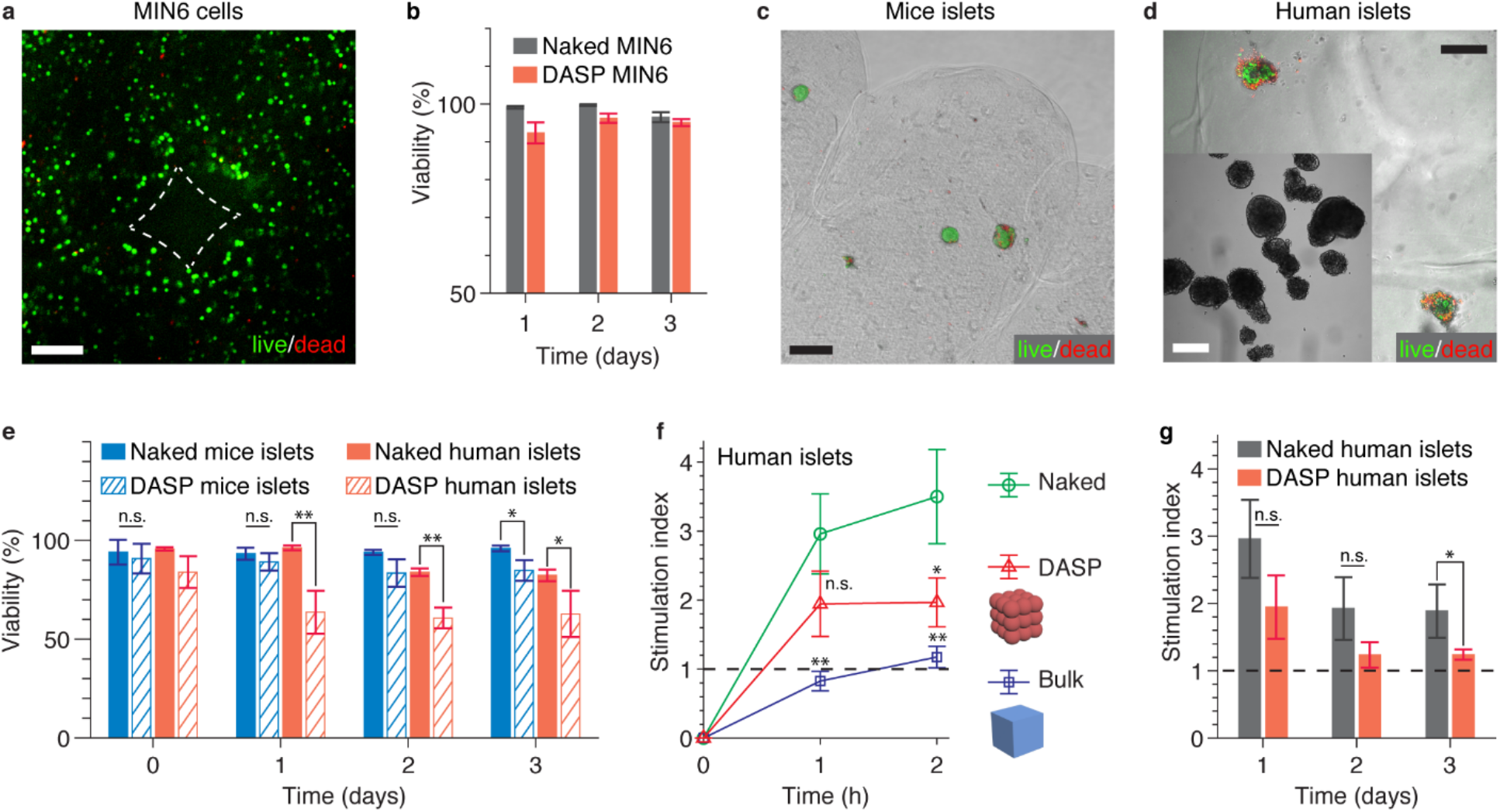
DASP scaffolds are cytocompatible and allow for fast, responsive, and sustained insulin release. a) A representative fluorescence confocal microscopy image from live/dead assay of DASP encapsulated MIN6 cells. The dashed lines outline the interparticle space as illustrated in **Figure 1d**. b) Viability of naked and DASP encapsulated MIN6 cells up to 3 days. c) Mice islets are encapsulated in a DASP printed lattice structure. The image is composed of three channels: the bright field that outlines the shape of the spherical hydrogel particles, the red fluorescence that corresponds to dead cells, and the green fluorescence that corresponds to live cells. d) Live/dead assay of DASP encapsulated human islets. Inset: bright-field image of human islets prior to encapsulation. e) Viability of naked and DASP encapsulated mice and human islets in 3 days. f) Glucose stimulation index from human islets in 2 hours after being stimulated by high glucose. Green circles: naked islets as positive control; red triangles: DASP encapsulated islets; blue squares: bulk hydrogel encapsulated islets as negative control. g) Glucose stimulation index of naked and DASP encapsulated human islets in 3 days. Error bar, standard deviation for *n=3*. \**p*<0.05, \*\**p*<0.01; *n*.*s*. not significant (Student’s t-test). Scale bars, 200 μm.

We further apply DASP to encapsulate human islets donated by patients (**SI Materials and Methods**). Unlike islets harvested from healthy mice, which often exhibit a smooth round shape (**Figure 7c**), clinical samples are often in unhealthy conditions such as long ischemia time, as reflected by the irregular shape and relatively loose protrusions of individual islets (inset of **Figure 7d**). As a result, clinical islets are often fragile and sensitive to mechanical shear. As expected, DASP encapsulated human islets exhibit significantly lower viability compared to their naked counterparts, as shown by the relatively large fraction of dead cells (**Figure 7d**, orange bars in **Figure e)**. Yet this difference becomes less significant by culturing the islets for longer time, as shown by the increased *p* value (**Figure 7e**). Nevertheless, the viability of DASP encapsulated human islets on day 3 remains at 62.8±11.7%, a rate sufficient for islet transplantation ^[37]^. These results suggest that DASP is suitable for encapsulating human pancreatic islets.

We monitor the insulin release of DASP encapsulated human islets and compare it with two control groups. One is a negative control in which human islets are encapsulated in a bulk bio-ink of the same composition and dimension, and the other is a positive control in which the naked cells are cultured in media (**SI Materials and Methods**). After being stimulated by glucose, within 1-hour DASP encapsulated human islets exhibit a glucose stimulation index of 2, comparable to that of the positive control (**Figure 7f**). Interestingly, the stimulation index of the bulk hydrogel, the negative control, is around 1, suggesting that the secreted insulin molecules, if any, are trapped in the bulk hydrogel. After culturing for 2 hours, the stimulation index increases slightly but remains low with the value around 1, suggesting that bulk bio-ink impairs the transport of molecules. By contrast, the stimulation index of DASP encapsulated islets is significantly higher with the value around 2. This is likely because, in a DASP printed lattice, the inter-particle space forms sub-millimeter channel to allow for efficient diffusion-based transport (outlined by dashed lines in **Figure 7a**). These results highlight the importance of the multiscale porous nature of DASP printed scaffolds, which enables fast, responsive insulin release.

The responsive insulin release of DASP printed scaffolds persists at longer times. Within 2 days, the stimulation index of DASP encapsulated islets is not significantly different from the naked counterparts (**Figure 7g**). Although the stimulation index becomes slightly lower on day 3, it is always larger than 1.2; this suggests a sustained responsive insulin release that meets the criteria for clinical transplantation ^[38,39]^. Collectively, our results show that DASP printed scaffolds allow fast, responsive, and sustained insulin release, critical to achieving synchronous and tight glucose level control.

## Conclusion

In summary, we have developed a 3D bioprinting technique that enables the digital assembly of spherical particles (DASP). Moreover, we have demonstrated the application of DASP in engineering mechanically robust multiscale porous scaffolds for encapsulating human islets for responsive insulin release, highlighting the potential of DASP in treating type 1 diabetes.

The novelty of DASP consists of three components: technology development, bio-ink design, and importantly, conceptual innovation. In the context of technology development, DASP is different from existing 3D printing techniques for manipulating spherical objects such as droplets and spheroids. For example, in embedded droplet printing, individual droplets are dispensed far apart from each other to form discrete patterns ^[17,40]^ or randomly stacked together to form thick structures ^[41]^. By contrast, DASP enables on-demand deposition of individual bio-ink droplets at prescribed locations to form integrated, organized structures. Moreover, DASP enables printing highly viscoelastic, non-Newtonian bio-inks in a cytocompatible environment; this contrasts with embedded droplet printing that relies on the interfacial tension between the immiscible aqueous and oil-like fluids to generate droplets, which, therefore, can handle low viscosity fluids only ^[17,40]^. Compared to bioprinting of spheroids ^[18,42]^ where cells are often subject to direct mechanical manipulation, in DASP cells are encapsulated in a hydrogel matrix, which avoids possible high mechanical shear-induced cell damage. Furthermore, the bio-ink particles can be assembled in a prescribed manner in a short time through controlled polymer swelling. This contrasts with the sophisticated surface chemistry such as lipid bilayers required to join droplets ^[33,43]^ and the slow cell migration required to fuse spheroids ^[18,42]^. Thus, DASP provides a facile, robust approach for the on-demand generation, deposition, and assembly of highly viscoelastic hydrogel particles in a bio-friendly environment.

In the context of biomaterials design, viscoelasticity and mesh size are perhaps two of the most important physical parameters ^[19,44]^. However, they are intrinsically coupled: stiffer, or more elastic, hydrogels are often of smaller mesh sizes. the concept of *hybrid* bio-inks provides a new way to independently control the viscoelasticity and the mesh size of hydrogels. The *hybrid* bio-ink is composed of two kinds of polymers interpenetrating with each other; one is an inert, long polymer that can form entanglements, whereas the other is a reactive polymer that can be crosslinked to form a network. Unlike typical approaches that use reversible bonds to tune the dynamics of hydrogels such as supramolecular networks ^[45,46]^, we exploit physical entanglements to control the viscoelasticity of bio-inks. During printing, the entanglements impart a relatively high viscosity and a high loss factor required for printing spherical bio-ink particles of good fidelity. After printing, the long polymers are trapped by the crosslinked network, but they can escape through reptation, leaving behind a structure with mesh size determined by the polymer concentration of the crosslinked hydrogel. The design concept is not limited to the alginate/PEO system in this study; instead, it should be general and will enable engineering bio-inks with independently controlled viscoelasticity and mesh size made of other polymers for tissue engineering ^[47–49]^.

Conceptually, DASP uses spherical hydrogel particles, not the conventional 1D filaments, as building blocks to create biological constructs. The spherical particles can be considered zero-dimensional voxels. When being used as building blocks, the location, composition, structure, and properties of a voxel can be locally defined to create complex 3D structures, as recently demonstrated by an origami structure and a millipede-like soft robot composed of soft and stiff elastic voxels ^[50]^. Thus, we believe that DASP has the potential to substantially expand the capability of extrusion bioprinting in engineering highly heterogeneous yet organized biological constructs ^[10]^.

## Supporting information

Movie S1_DASP printing process

Movie S2_DASP structure translation

## Supporting Information

Supporting Information is available from the Wiley Online Library or from the author.

## Acknowledgement

We thank Christopher Highley (University of Virginia) for critical reading of the manuscript.

## Author contributions

L.H.C. conceived the research. L.H.C., J.Z. and Y.H. designed the research. J.Z. performed the research including hardware modification, materials processing, physical and biological characterization, and data analysis. Y.H. performed the animal surgery, biological characterization, and data analysis. D.C. provided the modification scheme for the 3D printer and L.K. helped with drawing the design of the extrusion module. Z.H. helped with cell culture. L.H.C., J.Z. and Y.H. wrote the paper. K.K., S.G. and L.N. helped with the writing. All authors reviewed and commented on the paper.

## Data and materials availability

All data needed to evaluate the conclusions in the paper are present in the paper and/or the Supporting Information. Additional data related to this paper may be requested from L.H.C.

## References

[1] S. V Murphy, A. Atala, Nat. Biotechnol. 2014, 32, 773.

[2] H.-W. Kang, S. J. Lee, I. K. Ko, C. Kengla, J. J. Yoo, A. Atala, Nat. Biotechnol. 2016, 34, 312.

[3] S. C. Ligon, R. Liska, J. Stampfl, M. Gurr, R. Mülhaupt, Chem. Rev. 2017, 117, 10212.

[4] S. Nian, J. Zhu, H. Zhang, Z. Gong, G. Freychet, M. Zhernenkov, B. Xu, L.-H. Cai, Chem. Mater. 2021, 33, 2436.

[5] F. P. W. Melchels, J. Feijen, D. W. Grijpma, Biomaterials 2010, 31, 6121.

[6] B. Derby, Annu. Rev. Mater. Res. 2010, 40, 395.

[7] R. L. Truby, J. A. Lewis, Nature 2016, 540, 371.

[8] M. A. Skylar-Scott, S. G. M. Uzel, L. L. Nam, J. H. Ahrens, R. L. Truby, S. Damaraju, J. A. Lewis, Sci. Adv. 2019, 5, eaaw2459.

[9] J. Groll, T. Boland, T. Blunk, J. A. Burdick, D. W. Cho, P. D. Dalton, B. Derby, G. Forgacs, Q. Li, V. A. Mironov, L. Moroni, M. Nakamura, W. Shu, S. Takeuchi, G. Vozzi, T. B. F. Woodfield, T. Xu, J. J. Yoo, J. Malda, Biofabrication 2016, 8, 013001.

[10] L. Moroni, J. A. Burdick, C. Highley, S. J. Lee, Y. Morimoto, S. Takeuchi, J. J. Yoo, Nat. Rev. Mater. 2018, 3, 21.

[11] J. T. Muth, D. M. Vogt, R. L. Truby, D. B. Kolesky, R. J. Wood, J. A. Lewis, Adv. Mater. 2014, 26, 6307.

[12] T. J. Hinton, Q. Jallerat, R. N. Palchesko, J. H. Park, M. S. Grodzicki, H.-J. J. Shue, M. H. Ramadan, A. R. Hudson, A. W. Feinberg, Sci. Adv. 2015, 1, e1500758.

[13] T. Bhattacharjee, S. M. Zehnder, K. G. Rowe, S. Jain, R. M. Nixon, W. G. Sawyer, T. E. Angelini, Sci. Adv. 2015, 1, 4.

[14] C. B. Highley, C. B. Rodell, J. A. Burdick, Adv. Mater. 2015, 27, 5075.

[15] Q. D. Nguyen, D. V. Boger, Annu. Rev. Fluid Mech. 1992, 24, 47.

[16] A. Lee, A. R. Hudson, D. J. Shiwarski, J. W. Tashman, T. J. Hinton, S. Yerneni, J. M. Bliley, P. G. Campbell, A. W. Feinberg, Science (80-.). 2019, 365, 482.

[17] A. Z. Nelson, B. Kundukad, W. K. Wong, S. A. Khan, P. S. Doyle, Proc. Natl. Acad. Sci. U. S. A. 2020, 117, 5671.

[18] A. C. Daly, M. D. Davidson, J. A. Burdick, Nat. Commun. 2021, 12, 1.

[19] J. Li, D. J. Mooney, Nat. Rev. Mater. 2016, 1, 1.

[20] J. Malda, J. Visser, F. P. Melchels, T. Jüngst, W. E. Hennink, W. J. A. Dhert, J. Groll, D. W. Hutmacher, Adv. Mater. 2013, 25, 5011.

[21] N. Paxton, W. Smolan, T. Böck, F. Melchels, J. Groll, T. Jungst, Biofabrication 2017, 9, 044107.

[22] J. H. Y. Chung, S. Naficy, Z. Yue, R. Kapsa, A. Quigley, S. E. Moulton, G. G. Wallace, Biomater. Sci. 2013, 1, 763.

[23] J. S. Miller, K. R. Stevens, M. T. Yang, B. M. Baker, D. H. T. Nguyen, D. M. Cohen, E. Toro, A. A. Chen, P. A. Galie, X. Yu, R. Chaturvedi, S. N. Bhatia, C. S. Chen, Nat. Mater. 2012, 11, 768.

[24] D. B. Kolesky, R. L. Truby, A. S. Gladman, T. A. Busbee, K. A. Homan, J. A. Lewis, Adv. Mater. 2014, 26, 3124.

[25] D. B. Kolesky, K. A. Homan, M. A. Skylar-Scott, J. A. Lewis, Proc. Natl. Acad. Sci. U. S. A. 2016, 113, 3179.

[26] B. Grigoryan, S. J. Paulsen, D. C. Corbett, D. W. Sazer, C. L. Fortin, A. J. Zaita, P. T. Greenfield, N. J. Calafat, J. P. Gounley, A. H. Ta, F. Johansson, A. Randles, J. E. Rosenkrantz, J. D. Louis-Rosenberg, P. A. Galie, K. R. Stevens, J. S. Miller, Science (80-.). 2019, 364, 458.

[27] F. Lim, A. M. Sun, Science (80-.). 1980, 21, 908.

[28] R. Satchi-Fainaro, R. Duncan, C. M. Barnes, in Adv. Polym. Sci., 2006, pp. 1–65.

[29] M. Rubinstein, R. H. Colby, Polymer Physics, Oxford University Press, Oxford, UK, 2003.

[30] K. Y. Lee, D. J. Mooney, Prog. Polym. Sci. 2012, 37, 106.

[31] H. Wen, J. Hao, S. K. Li, J. Pharm. Sci. 2013, 102, 892.

[32] T. Desai, L. D. Shea, Nat. Rev. Drug Discov. 2017, 16, 338.

[33] G. Villar, A. D. Graham, H. Bayley, Science (80-.). 2013, 340, 48.

[34] G. Villar, A. J. Heron, H. Bayley, Nat. Nanotechnol. 2011, 6, 803.

[35] A. J. Vegas, O. Veiseh, M. Gürtler, J. R. Millman, F. W. Pagliuca, A. R. Bader, J. C. Doloff, J. Li, M. Chen, K. Olejnik, H. H. Tam, S. Jhunjhunwala, E. Langan, S. Aresta-Dasilva, S. Gandham, J. J. McGarrigle, M. A. Bochenek, J. Hollister-Lock, J. Oberholzer, D. L. Greiner, G. C. Weir, D. A. Melton, R. Langer, D. G. Anderson, Nat. Med. 2016, 22, 306.

[36] Q. Liu, A. Chiu, L. H. Wang, D. An, M. Zhong, A. M. Smink, B. J. de Haan, P. de Vos, K. Keane, A. Vegge, E. Y. Chen, W. Song, W. F. Liu, J. Flanders, C. Rescan, L. G. Grunnet, X. Wang, M. Ma, Nat. Commun. 2019, 10, 1.

[37] C. Ricordi, J. S. Goldstein, A. N. Balamurugan, G. L. Szot, T. Kin, C. Liu, C. W. Czarniecki, B. Barbaro, N. D. Bridges, J. Cano, W. R. Clarke, T. L. Eggerman, L. G. Hunsicker, D. B. Kaufman, A. Khan, D. E. Lafontant, E. Linetsky, X. Luo, J. F. Markmann, A. Naji, O. Korsgren, J. Oberholzer, N. A. Turgeon, D. Brandhorst, A. S. Friberg, J. Lei, L. J. Wang, J. J. Wilhelm, J. Willits, X. Zhang, B. J. Hering, A. M. Posselt, P. G. Stock, A. M. J. Shapiro, Diabetes 2016, 65, 3418.

[38] K. K. Papas, T. M. Suszynski, C. K. Colton, Curr. Opin. Organ Transplant. 2009, 14, 674.

[39] A. Pileggi, C. Ricordi, N. S. Kenyon, T. Froud, D. A. Baidal, A. Kahn, G. Selvaggi, R. Alejandro, in Clin. Transpl., 2004, pp. 177–204.

[40] L. Cai, J. Marthelot, P. T. Brun, Proc. Natl. Acad. Sci. U. S. A. 2019, 116, 22966.

[41] H. J. Mea, L. Delgadillo, J. Wan, Proc. Natl. Acad. Sci. U. S. A. 2020, 117, 14790.

[42] B. Ayan, D. N. Heo, Z. Zhang, M. Dey, A. Povilianskas, C. Drapaca, I. T. Ozbolat, Sci. Adv. 2020, 6, 1.

[43] F. G. Downs, D. J. Lunn, M. J. Booth, J. B. Sauer, W. J. Ramsay, R. G. Klemperer, C. J. Hawker, H. Bayley, Nat. Chem. 2020, 12, 363.

[44] O. Chaudhuri, L. Gu, D. Klumpers, M. Darnell, S. A. Bencherif, J. C. Weaver, N. Huebsch, H. P. Lee, E. Lippens, G. N. Duda, D. J. Mooney, Nat. Mater. 2016, 15, 326.

[45] X. Du, J. Zhou, J. Shi, B. Xu, Chem. Rev. 2015, 115, 13165.

[46] E. A. Appel, J. del Barrio, X. J. Loh, O. A. Scherman, Chem. Soc. Rev. 2012, 41, 6195.

[47] M. W. Tibbitt, K. S. Anseth, Biotechnol. Bioeng. 2009, 103, 655.

[48] J. L. Drury, D. J. Mooney, Biomaterials 2003, 24, 4337.

[49] J. A. Burdick, G. D. Prestwich, Adv. Mater. 2011, 23, 41.

[50] M. A. Skylar-Scott, J. Mueller, C. W. Visser, J. A. Lewis, Nature 2019, 575, 330.

