## supporting information for "Voxelated Bioprinting of Mechanically Robust Multiscale Porous Scaffolds for Pancreatic Islets"

**This PDF file includes**

- SI Materials and Methods
- Figure S1. Setup of DASP.
- Figure S2. Crosslinking kinetics of alginate hydrogel.
- Figure S3. Swelling kinetics of an alginate hydrogel particle.
- Figure S4. Bio-inks are cytocompatible.
- Movie S1. Implementation of DASP.
- Movie S2. Manipulating a DASP printed structure.

### SI Materials and Methods

**Materials and reagents**. Gelatin from porcine skin (gel strength 300, Type A, Cat. No. G2500), alginic acid sodium salt from brown algae (medium viscosity, Cat. No. A2033), poly(ethylene oxide) (PEO) with MW of 5000, 1000, 300, 100 kDa (Cat. No. 189472, 182001, 372781, 181986), and poly(ethylene glycol) (PEG) with MW of 35 kDa (Cat. No.81310) were purchased from Sigma-Aldrich (USA). Fluorescent labeling reagents including EDC (Cat. No. E7750), Sulfo-NHS (Cat. No. 56485) and fluoresceinamine (Cat. No. 201626) were purchased from Sigma-Aldrich (USA). Fluorescently labeled dextran (Texas Red™, MW=70,000 Da, ex 595/em 615, Cat. No. D1830) was purchased from Fisher Scientific (USA). Minimum Essential Medium (MEM, low glucose, without calcium, Cat. No. 11380-037), Dulbecco's Modified Eagle Medium (DMEM, high glucose, GlutaMax, Cat. No. 10566-016), DMEM (high glucose, without calcium, Cat. No. 21068-028), DMEM (without glucose, glutamine, phenol red, and sodium pyruvate, Cat. No. A14430-01), Roswell Park Memorial Institute 1640 Medium (RPMI 1640, Cat. No. 72400-047), FBS (Cat. No. 10438-018), Pen/Strep (Cat. No. 15140-122), 2-Mercaptoethanol (Cat. No. 21985-023), Dulbecco's phosphate buffered saline (DPBS, without calcium, Cat. No. MT21031CM), Hank's balanced salt solution (HBSS, without calcium, Cat. No. 14175-079) were purchased from Gibco, Fisher Scientific (USA). Connaught Medical Research Laboratories 1066 medium (CMRL 1066, Cat. No. 99-663-CV) was purchased from Corning (USA). Chemical dye for live/dead assay including fluorescein diacetate (Cat. No. F7378), propidium iodide (Cat. No. P4170) were purchased from Sigma-Aldrich (USA).

**MIN6 cells.** MIN6 cells are cultured in DMEM culture media (4.5 g/L D-Glucose) supplemented with 10% FBS, 1% Pen/Strep, 0.0005% 2-Mercaptoethanol and 20 mM HEPES.

**Mice islets.** Under the approval by UVA Institutional Animal Care and Use Committee (IACUC) (No. 4196-10-20), we isolate mice islets from the pancreata of 10-12 week-old male and female mice (C57BL/6J, Jackson Laboratory, ME) following the standard procedure [1]. In brief, the pancreata are injected with 0.375 mg/mL of Collagenase P (Roche Diagnostics, IN) and then digested at 37 °C for 12 min. The digested islets are purified using Ficoll-Hypaque density gradient centrifugation, a method that isolates islets based on their difference in density compared to other elements in the islet suspension [1]. The purified islets are cultured in RPMI 1640 medium supplemented with 10% FBS and 1% Pen/Strep at 37°C under 5% $CO_2$.

**Human islets.** Human pancreatic islets are isolated in Good Manufacturing Practice (GMP) facility at University of Virginia, following a published protocol approved by UVA, which is a modified method based on a published procedure [2,3]. The pancreata are obtained from a donor (Male, Caucasian, age 48, body mass index 29.7, non-diabetic) through Organ Procurement Organizations (OPO) with research consent. In brief, the pancreata are trimmed and distended using Liberase HI (Roche Applied Science, IN), and digested in a modified Ricordi chamber. After collection and washing steps, the digested tissue is further purified using a cell separator (Cobe 2991, Cobe Inc., CO) following University of Illinois at Chicago UW/Biocoll method (UIC-UB), and then cultured in CMRL 1066 medium supplemented with 5% human albumin (CSL Behring LLC., IL) at 37°C under 5% $CO_2$.

**Hardware of DASP.** The hardware of DASP includes a 3D motion system and a customized printing nozzle. The 3D motion system consists of a XYZ stage and an additional motorized vertical axis; the former allows positioning the printing nozzle in space, whereas the latter allows independent control over the extrusion speed of the printing nozzle. We build the 3D motion system by replacing the hotmelt extruder of a desktop printer (JGURORA z-603s) with a custom-made extrusion module (Figure S1a). The extrusion module is built based on a linear screw (T8) actuator which converts the rotary motion of a stepper motor (NEMA 17) into linear motion. We use the extrusion module to drive a glass syringe (1 mL, Shanghai Bolige Industrial & Trade Co., Ltd) with a speed controlled by G-code (Figure S1b).

To fabricate the printing nozzle, we use a micropipette puller (P-1000, Sutter Instrument, Inc.) to taper a cylindrical glass capillary (World Precision Instruments, Inc.) of inner and outer diameters 0.58 mm and 1.00 mm, respectively, to a diameter of 20 μm and then carefully sand it to a desired diameter. We sleeve the glass capillary to a dispensing needle (26G, McMaster-Carr), and seal the interstitial space between the two using 5-Minute Epoxy (Devcon) (Figure S1c-e). We mount the glass capillary sleeved needle to a glass syringe, which can be driven by the extrusion module.

**Gelatin supporting matrix.** To prepare the supporting matrix, we first make a gelatin hydrogel and then fragment the hydrogel into microparticles. Specifically, we dissolve gelatin in a calcium solution (6 mM calcium chloride, 140 mM sodium chloride) with a concentration of 1.5% (w/v) at an alleviated temperature 50 ℃, and then cool the solution at 4 ℃ for 12 hours to solidify the solution. In a typical procedure, we mix 200 mL hydrogel with 30 mL calcium solution (6 mM calcium chloride, 140 mM sodium chloride), and use a blender at 4000 rpm for 40 sec to break the hydrogel into microparticles. The mixture is centrifuged at 2000 rpm to remove the supernatant of foam then degassed using house vacuum for 2 min to remove bubbles. This results in a gelatin slurry that is an optically semi-transparent and mechanically yield-stress fluid that can self-heal in less than a second.

**Bio-inks.** To prepare the alginate-based bio-inks used for printability study, we dissolve alginic acid sodium salt in DI water at 4.0% (w/v), and then sonicate the mixture for 2 h at 60 °C to make a homogenous solution. All *hybrid* bio-inks are prepared by dissolving alginate and PEO sequentially in DI water at 60 °C. All bio-inks are stored in a 4 °C refrigerator before use. For cell and islets encapsulation, the DI water is replaced by cell culture medium, as described in individual sections below.

**Fluorescently labeled alginate.** We use a previously developed method to fluorescently label alginate [4]. In brief, we dissolve alginate in PBS at a concentration of 0.5% (w/v), equivalent to 230 mM carboxylic groups, and add EDC and Sulfo-NHS with a final concentration of 23 mM for each. We stir the solution at room temperature for 2 h, add fluoresceinamine with a final concentration of 45 mM, and then continue the stirring for 48 hours. Afterward, we transfer the solution to dialysis bags (Spectra/Por, VWR International, molecular weight cut off, 6-8 kDa, Cat. No. 28170-138) and dialyze the solution against firstly 3.5 L DI water (refreshed every 6 hours, 4 times), secondly DI water with 1M NaCl (refreshed every 6 hours, 4 shifts), and finally DI water (refreshed every 6 hours, 4 times). This process completely removes the unreacted EDC, Sulfo-NHS, and fluoresceinamine. The dialyzed alginate solution is lyophilized to obtain a dried alginate and stored in a 4 °C refrigerator for further usage. All operations are protected from the light.

**Rheometry.** Rheological measurements are performed using a stress-controlled rheometer (Anton Paar MCR 302) equipped with parallel-plate geometries of diameter 25 mm for bio-inks and 8 mm for crosslinked hydrogels at 20℃. To characterize the yield-stress behavior of the supporting matrix, we conduct a stress sweep from 0.1 to 1000 Pa at an oscillatory frequency of 1 Hz. To characterize the self-healing behavior, we monitor the viscoelastic properties of the supporting matrix subjected to a periodic destructive high shear strain. For each cycle, we increase the shear strain to 1000% within 1 sec, and then apply an oscillatory shear strain of 1% at a frequency of 1 Hz for 200 seconds, during which both $G'$ and $G''$ are measured.

To determine the gelation time of the alginate-based bio-inks, we monitor in *real-time* the $G'$ and $G''$ at an oscillatory shear frequency of 1 Hz and a shear strain of 1%. We choose a gap size of 1 mm, comparable to the dimension of a printed particle. Once the reading is stable, we apply an excess amount of calcium contained gelatin matrix to fully cover the peripheral of the geometry. Simultaneously, $G'$ and $G''$ exhibit a sharp increase associated with the instantaneous physical contact between the matrix and the geometry. The effects of this physical contact on measured moduli are corrected by referencing to the measurements using the supporting matrix containing no calcium ions.

We measure the dependence of viscosity on shear rate $\dot{\gamma}$ in the range of 0.01 to 500 1/sec. For all bio-inks suitable for DASP, the viscosity is nearly constant at shear rates (<0.1 $s^{-1}$) but decreases rapidly at $\dot{\gamma} > 1\ s^{-1}$ and becomes nearly two orders of smaller at high shear rates (>100 $s^{-1}$). These results suggest that bio-inks suitable for DASP are shear-thinning fluids (Figure 4b). We take the value at the lowest shear rate 0.01 $s^{-1}$ as the viscosity of a bio-ink.

To characterize the loss factor of a bio-ink, we measure the dependencies of $G'$ and $G''$ on strain in the range of 0.1% to 10000% at an oscillatory frequency of 1 Hz. As expected for shear-thinning fluids, both $G'$ and $G''$ are nearly constant at shear strain lower than 50% but decrease dramatically above ~200% strain (Figure 4c). We take the values $G'$ and $G''$ at 0.1% strain to determine the loss factor.

To characterize the shear moduli of hydrogels made from alginate and *hybrid* alginate/PEO bio-inks, we apply an oscillatory shear with a frequency in the range of 0.1 to 100 Hz with at a strain of 0.5 %. To prepare a hydrogel sample, we use the lid of a cell culture dish (Greiner Bio-One CELLSTAR™, Fisher Scientific Cat. No. 07-000-584) to mold a bio-ink film with the thickness of approximately 1 mm. The film is solidified by immersing it into a crosslinking solution, DI water with 50 mM $Ca^{2+}$. Then, we transfer the crosslinked film to DMEM (without glucose, glutamine, phenol red, and sodium pyruvate) and incubate for 30 min to fully equilibrate the film with the medium. To match the sample size with that of the rheometer geometry, we use a hole puncher to trim the film to obtain a disk with 8 mm in diameter. We load the sample to the bottom geometry of the rheometer, and lower the upper geometry to contact with the sample, as indicated by a positive normal force around 0.1 N.

**Characterization of bio-ink mesh size.** We use two independent methods to characterize the bio-ink mesh size. For the first method, we measure the shear modulus $G$ of the crosslinked bio-ink hydrogel. Using the relation that the network shear modulus is $k_B T$ per volume occupied by a network strand, we calculate the mesh size by $\xi \approx (k_B T/G)^{1/3}$.

For the second method, we encapsulate protein mimics, a fluorescently labeled dextran (Texas Red, 70 kDa), within a hydrogel particle and quantify release profile of dextran molecules. Specifically, we dissolve in a bio-ink the fluorescently labeled dextran at a concentration of 2 mg/mL. A bio-ink droplet is deposited into a crosslinking solution, DI water with 50 mM $Ca^{2+}$ to solidify the particle. We immediately transfer the crosslinked particles to DMEM (without glucose, glutamine, phenol red, and sodium pyruvate) and incubate the particles for 30 min to fully equilibrate them with the medium. Importantly, the solutions for crosslinking and washing the bio-ink droplets contain the same 2 mg/mL dextran as that in the bio-ink droplet. This prevents the leak of encapsulated dextran molecules. We then replace the washing medium by a fresh DMEM medium that contains no dextran, during which the fluorescence of the particle is monitored using confocal laser scanning fluorescence microscopy (Leica SP8). We use the half-decay time of the fluorescence intensity to assess the mesh size of the hydrogel particle.

**Printing and characterization of bio-ink particles and lattice structures.** To print a hydrogel particle, we use a 'forward-then-backward' mode to drive the glass capillary in the supporting matrix. Specifically, we position the printing nozzle at a desired location, extrude a prescribed amount of bio-ink, move the printing nozzle towards the next position by 3.5 mm, and then move the nozzle back to the targeted position (Figure S2f). Once the printing is completed, the printed feature is kept in the supporting matrix for 3 minutes. Afterwards, we immerse the supporting matrix in a bath of 50 mM calcium solution (75 mM NaCl) at 37 ℃ for 5 min (Movie. S1). This not only dissociate the gelatin supporting matrix but also completely crosslinks the alginate hydrogels.

To visualize the shape of individual bio-ink particles, we use fluorescently labeled alginate to prepare all bio-ink formulations. Using the same printing conditions as described above, we deposit a bio-ink droplet at a location 2 mm below the surface of the supporting matrix. This distance is short enough to allow us to use a digital microscope camera (Hayear, 16 MP) to characterize the morphology of the droplet.

The printed 3D structures are imaged after the removal of the supporting matrix. To visualize the junctions between neighboring particles, we immerse a structure in DI water that contains a red food dye (McCormick, red), which preferentially aggregates at the boundary between particles, as shown Figure 6b.

**Compression tests.** Because the bio-inks are soft, the force required to deform the material is small. To this end, we use a rheometer (Anton Paar MCR 302) with a normal force resolution of 0.5 mN to perform the compression tests. The sample, in the form of either a bulk material or a printed 3D lattice, is placed onto the bottom geometry. We lower the upper geometry to contact with the sample, at which the normal force is slightly above zero. During the compression measurements, the moving profile of the upper plate is pre-setup to exert large compression at a fixed strain rate 0.005/sec. We record the normal force, gap size, and time, and calculate the stress and strain based on the pre-measured dimensions of the samples.

**DASP printing scaffolds encapsulated with MIN6 cells.** To obtain relatively large amount of MIN6 cells, we culture them in multiple T75 flasks. To harvest the cells, we rinse each flask using 1x PBS, apply 3 mL Trypsin-EDTA for 3 min to detach the cells from the substrate, and then add 6 mL DMEM culture media to neutralize the Trypsin-EDTA. We collect the suspension of cells and centrifugate it at 600 g for 5 min to obtain a pellet of cells, which are subsequently suspended in DMEM (high glucose, without calcium) to reach a concentration of 15 million/mL.

To prepare alginate-based bio-inks, we dissolve sterilized alginate in DMEM (high glucose, without calcium) with a concentration of 5.0% (w/v). For *hybrid* alginate/PEO bio-inks, the concentration of PEO is relatively high, which generates osmotic pressure that impairs cell viability. To this end, we dissolve alginate and PEO in a reduced DMEM, which consists of 1:1

volume ratio of DMEM and mannitol solution, to alleviate the osmolarity to that of the physiological condition. We then mix the bio-ink with the cell suspension at 4:1 volume ratio to reach the final concentration of 4.0% (w/v) for alginate and 3 million/mL for MIN6 cells. Using the printing and washing processes described above, we DASP print MIN6 cells encapsulated scaffolds and transfer them to MIN6 culture medium for subsequent cytocompatibility test.

**DASP printing 3D scaffolds encapsulated with mice and human islets.** Using pipette under a stereoscope, we hand pick a desired number of islets, which are further concentrated by centrifuging at 300 g for 3 min to obtain a pellet of islets. The pellet is resuspended in MEM medium (low glucose, without calcium) to reach a concentration of 5000 islets/mL. To prepare the bio-ink, we dissolve sterilized alginate in MEM with a concentration of 5% w/v, and then mix the solution with the islet suspension at 4:1 volume ratio to reach the final concentration of 4% w/v for alginate and 1000 islets/mL for islets. To accommodate the size of islets, 150 µm and 300 µm in diameter respectively for mice and human islets, we use stainless-steel dispensing needles of 26G (ID 250 µm) and 25G (ID 300 µm) respectively as the printing nozzle. Using the printing and washing processes described above, we DASP print scaffolds encapsulated with islets and transfer the scaffolds to culture medium (RPMI 1640 for mice islets and CMRL 1066 for human islets) for subsequent cytocompatibility and insulin release assays.

**Cytocompatibility of DASP.** We use live/dead assay to characterize the cytocompatibility of DASP. To prepare the dye solution for the assay, we add the stock propidium iodide solution, 750 µM in DPBS, and fluorescein diacetate solution, 1 mM in DMSO, to HBSS without calcium to respectively reach final concentrations of 3.75 µM and 0.2 µM. To stain naked MIN6 cells, mice and human islets, as well as the encapsulated ones, we remove the culture media and add the dye solution at 0.5 mL/well for 48-well plate and 1 mL/well for 24-well plate. Using a 10x dry objective with a numerical aperture of 0.3 on a confocal microscope (Leica SP8) equipped with a 37 ℃ incubator, we acquire bright field and fluorescence images simultaneously using multitrack mode. Specifically, we use HeNe laser (552 nm) as the excitation source for propidium iodide and Argon laser (488 nm) as the excitation source for fluorescein diacetate. The fluorescence emission is collected by PMT detectors through filters with bandpass 600/700 nm for propidium iodide and

500/540 nm for fluorescein diacetate. Similar methods are used to characterize the viability of mice and human islets.

The encapsulated cells are not on the same focus plane; instead, they are randomly distributed in hydrogel particles. To quantify live/dead cells, we use fluorescence confocal microscopy to acquire a stack of images along $z$-axis to capture all cells, and construct a $z$-projection image by summing up all slices; examples of such $z$-projection images are shown in Figure 7c, d. This allows us to quantify the fraction of live cells, which is defined as the ratio between the number of green pixels to the total number of green and red pixels.

**Glucose-stimulated insulin secretion (GSIS) test.** We use Krebs Ringer buffer (KRB) as the medium for GSIS test, which consists of three steps: washing, equilibration, and stimulation. Before starting the test, we revive a sample, in the form of naked, DASP encapsulated, or bulk hydrogel encapsulated islets, by pre-culturing it in a well of 24-well plate for 24 hours. In the washing step, we rinse the sample using 1 mL basal-glucose solution (KRB supplemented with 2 mM D-glucose) for twice with the interval of 1 hour. Such a washing process removes the residue insulin secreted during the pre-culturing period. In the equilibration step, we exchange the basal medium with 1 mL fresh basal-glucose solution and collect all the medium after 1 hour's incubation. In the stimulation step, we add 1 mL high-glucose solution (KRB supplemented with 25 mM D-glucose), incubate for 1 hour, and then collect all the medium. Parallel samples are used to collect the medium at different time points.

Compared to encapsulated islets, naked ones are free to move, resulting in a practical challenge in exchanging and collecting medium without losing islets. To this end, we culture naked islets in a transwell insert with the pore size of 8 μm (Falcon, Fisher Scientific, Cat. No. 08-771-21); this insert separates the islets from the media in the well yet is permeable to biomolecules. In a typical washing step, for example, we respectively add 700 μL and 300 μL medium in the well and the insert, incubate for a desired time, and exchange the medium. The 700 μL is replaced by a fresh medium using pipette. For the transwell insert, we gently dip its bottom onto a sterilized Kimwipe (Kimtech, VWR International, Cat. No. 470224-038), which generates capillary force to completely remove the media on the apical side of the transwell, and then add 300 μL fresh medium. Similar procedures are applied to the equilibration and stimulation steps.

To quantify insulin release, we centrifuge the collected medium obtained from the stimulation phase at 300 g for 3 min to remove possible cells and debris. We aliquot 200 μL supernatant and measure its insulin concentration $c_{ins}$ using Human Insulin ELISA kits (ALPCO Ltd, Fisher Scientific, Cat. No. 50-751-3654). The stimulation index is calculated by normalizing the insulin concentration upon stimulation to the basal value: $Stimulation\ index = c_{ins}^{stim}/c_{ins}^{basal}$.

**Statistical analysis.** All data are shown as mean ± S.D. with sample size *n*=3. Comparisons among all groups are evaluated using Student's t-test. *$p$<0.05, **$p$<0.01, ***$p$<0.001, ****$p$<0.0001, and $p$>0.05: no significant difference.

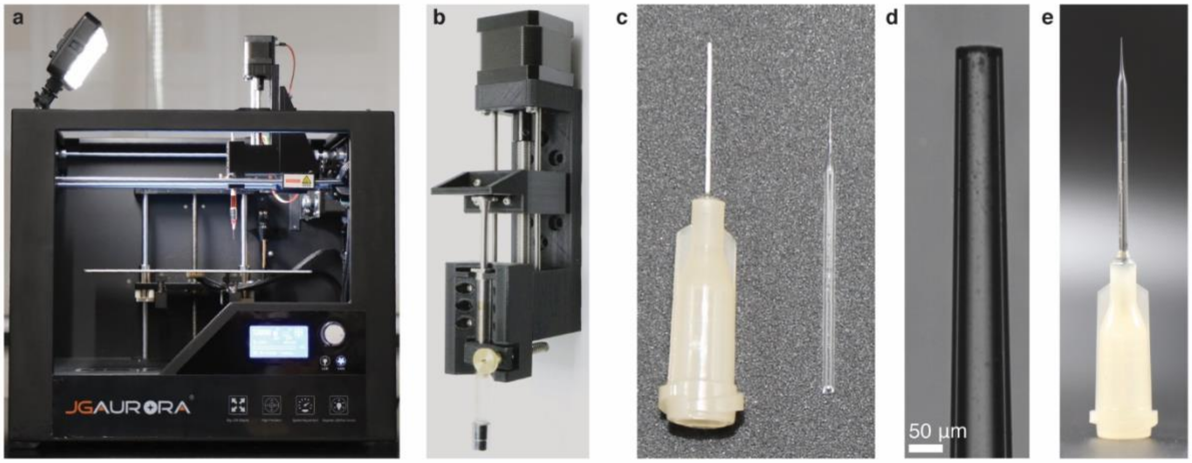


**Figure S1. Setup of DASP.** a) A photograph of the 3D motion system with a computer-controlled syringe pump. b) A zoom-in photo of the computer-controlled syringe pump. c) Parts of the printing nozzle: left, a 26G metal dispensing needle with a Luer-lock connection; right, a tapered cylindrical glass capillary with the body diameter of 1 mm and the tip diameter of 80 μm. d) An optical image for the tip of the printing nozzle. e) The glass capillary is sleeved to the dispensing needle and sealed with epoxy.

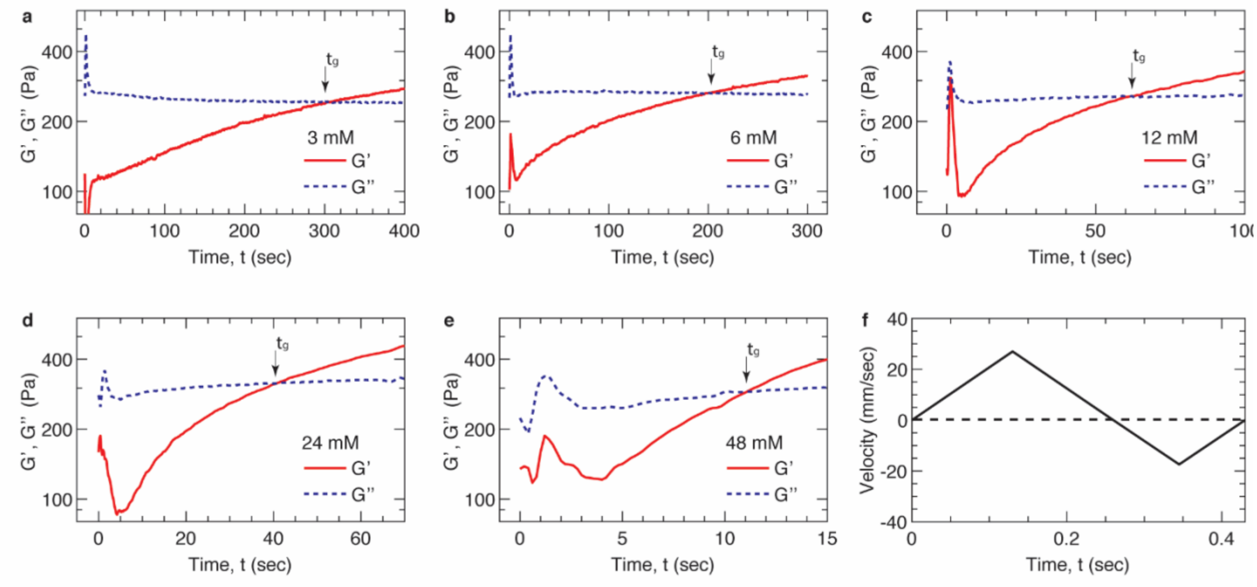


**Figure S2. Crosslinking kinetics of alginate hydrogel**. a-e) *Real-time* characterization of storage ($G'$, solid lines) and loss ($G''$, dashed lines) moduli for the alginate solution in the presence calcium ions at the concentrations of 3, 6, 12, 24, 48 mM. The peaks for $G'$ and $G''$ are associated with the instantaneous friction between the oscillatory geometry and the gelatin supporting matrix (see **Materials and Methods**). f) The velocity profile of the ‘forward-then-backward’ mode used to move the printhead with the distance of 1 mm.

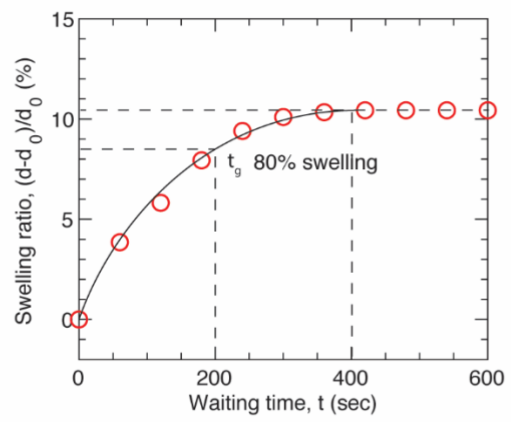


**Figure S3. Swelling kinetics of an alginate hydrogel particle.** A fluorescently labelled hydrogel particle made of $Alg_{4.0}$ is deposited at a location 2 mm below the surface of the supporting matrix. The swelling process is visualized under a digital microscope camera to monitor the diameter of the droplet. Swelling ratio is defined as $100\% \times (d - d_0)/d_0$, in which $d_0$ and $d$ are respectively the initial and *real-time* particle diameter. At about 600 seconds, the particle reaches equilibrium swelling with the diameter increased by nearly 11%. The particle reaches 80% of the equilibrium swelling at about 200 seconds, comparable to the gelation time $t_g$ of alginate (**Figure S2b**). Solid line represents the guidance for eye.

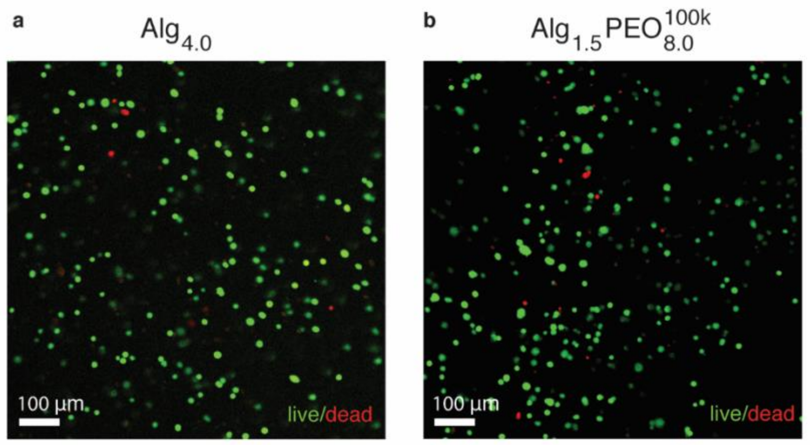


**Figure S4. Bio-inks are cytocompatible.** Representative laser scanning fluorescence confocal microscopy images from live/dead assay of encapsulated MIN6 cells in hydrogel made from a) 4.0% (w/v) alginate and b) a hybrid bio-ink consisting of 1.5% (w/v) alginate and 8.0% (w/v) PEO (MW 100 kDa).

**Movie S1 (.mp4). Implementation of DASP.**

**Movie S2 (.mp4). Manipulating a DASP printed structure.**

### References


[1] J. G. Avila, Y. Wang, B. Barbaro, A. Gangemi, M. Qi, J. Kuechle, N. Doubleday, M. Doubleday, T. Churchill, P. Salehi, J. Shapiro, L. H. Philipson, E. Benedetti, J. R. T. Lakey, J. Oberholzer, *Am. J. Transplant.* **2006**, *6*, 2861.

[2] M. Qi, B. Barbaro, S. Wang, Y. Wang, M. Hansen, J. Oberholzer, *J. Vis. Exp.* **2009**, 37.

[3] M. Qi, B. Barbaro, S. Wang, Y. Wang, M. Hansen, J. Oberholzer, *J. Vis. Exp.* **2009**, 3.

[4] B. L. Strand, Y. A. Mørch, T. Espevik, G. Skjåk-Bræk, *Biotechnol. Bioeng.* **2003**, *82*, 386.
